# Construction of an exome-wide risk score for schizophrenia based on a weighted burden test

**DOI:** 10.1101/145961

**Authors:** David Curtis

## Abstract

Polygenic risk scores obtained as a weighted sum of associated variants can be used to explore association in additional data sets and to assign risk scores to individuals. The methods used to derive polygenic risk scores from common SNPs are not suitable for variants detected in whole exome sequencing studies. Rare variants which may have major effects are seen too infrequently to judge whether they are associated and may not be shared between training and test subjects. A method is proposed whereby variants are weighted according to their frequency, their annotations and to the genes they affect. A weighted sum across all variants provides an individual risk score. Scores constructed in this way are used in a weighted burden test and are shown to be significantly different between schizophrenia cases and controls using a five-way cross validation procedure. This approach represents a first attempt to summarise exome sequence variation into a summary risk score, which could be combined with risk scores from common variants and from environmental factors. It is hoped that the method could be developed further.

## Introduction

Polygenic scores have found widespread application since they were used in a GWAS of schizophrenia (Purcell et al. 2009). In this study, very large numbers of variants showing weak association signals in a training set of cases with schizophrenia and controls were used to produce a score which was higher in a test set of cases with schizophrenia than controls and was also in increased in subjects with bipolar disorder. As noted previously, the two main functions of the polygenic score are to demonstrate that variants selected from the training set are associated with the trait in the test set and to provide an overall assessment of an individual’s genetic risk (Dudbridge 2013). The score consists of a weighted sum of the scores of the variant alleles possessed by the test subject. Different methods can be used to select the variants to be included and to assign their weights (Euesden et al. 2015). A key feature is that a large number variants is used and it is understood that many will not in fact be truly associated with the trait. As the sample size for the training set increases, so does the power to distinguish the truly associated variants and hence the polygenic score can become a more accurate determinant of genetic risk.

Recently, association studies have been carried out involving whole exome sequencing of thousands of subjects. For non-Mendelian diseases it is expected that there will be contributions to genetic risk from a number of different loci but it is not straightforward to obtain polygenic scores using a process similar to that which is appropriate for GWAS SNPs. There are several of reasons for this. One is that exome sequencing detects a very large number of variants and that rare variants tend to have weaker LD relationships than common SNPs, meaning that the number of independent signals is greater. However the main problem is that very rare variants may have major effects on risk but that they are so infrequent that there is very little information as to which variants are individually associated. A recent exome-sequencing study of schizophrenia concluded that singleton variants, observed only in one study subject and never in ExAC, did have major effects (Genovese et al. 2016). One could never hope to derive a polygenic score using such variants because one could never know if a specific singleton variant had an effect or not and even if it did one would not expect to see it in a test subject. Another difference between GWAS SNPs and exome sequence variants is that the latter have a higher intrinsic information content. Some GWAS SNPs may be identified as being associated with gene expression but for many SNPs one can make only weak inferences about likely effect and one may not even know which gene is functionally relevant. However, an exome variant can be annotated and one can make reasonable predictions about which gene is likely to be affected and the nature of the effect. It would be desirable to incorporate such information into a score designed to reflect genetic risk.

Overall, it seems that a polygenic risk score derived from exome sequence variants should be able to utilise variants which have not been seen in the training set but which are, in some defined way, similar to them. It will be expected that the risk score will make use of information about the likely effect of the variant and about the gene or type of gene which it affects. Such a scheme was devised and applied to the schizophrenia case-control dataset.

## Methods

The data analysed consisted of whole exome sequence variants downloaded from dbGaP from a Swedish schizophrenia association study containing 4968 cases and 6245 controls (Genovese et al. 2016). The original analysis demonstrated that there was an excess of damaging ultra-rare variants among cases, concentrated in particular gene sets. This sample included the 2545 cases and 2545 controls used for previously reported exome sequence association studies (Purcell et al. 2014; Curtis 2013; Curtis 2016). The dataset was managed and annotated using the GENEVARASSOC program which accompanies SCOREASSOC (https://github.com/davenomiddlenamecurtis/geneVarAssoc). Version hg19 of the reference human genome sequence and RefSeq genes were used to select variants on a gene-wise basis.

A number of QC processes were applied. Variants were excluded if they did not have a PASS in the information field and individual genotype calls were excluded if they had a quality score less than 30. Variants were also excluded if there were more than 10% of genotypes missing or of low quality in either cases or controls or if the heterozygote count was smaller than both homozygote counts in both cohorts. A preliminary weighted burden test analysis using variants with MAF<0.01 was carried out using SCOREASSOC (Curtis 2012). This identified several genes which had a significant excess of rare, functional variants in cases but on closer examination it emerged that these results were driven by variants which were reported in ExAC to have a markedly different allele frequency in Finnish as opposed to non-Finnish Europeans (Lek et al. 2015). In order to address this issue we set out to identify those subjects who appeared to have a substantial Finnish component to their ancestry. To do this, for each subject the genotype of the variant with the highest MAF in each of 18349 genes was used to calculate an odds ratio based on the Finnish versus non-Finnish European allele frequencies presented in ExAC r.03. The logs of these odds ratios were then summed to produce a measure denoted as the F score. The distributions of the F scores in cases and controls were plotted and each distribution was mostly normally distributed but had an extended right tail, indicating that a proportion of both cases and controls were likely to have substantial Finnish ancestry. The right tail was larger in the cases and overall the cases had significantly higher F scores than controls (t=16.4, df=11212, p=2.2e-16). Overall the mean F score was -13.0 with SD 24.3. A cut-off value of 10 was chosen to exclude the right tails, and subjects with a higher score were removed, comprising 743 cases and 411 controls. When the gene-wise weighted burden tests were repeated on the reduced sample of 4225 cases and 5834 controls the previous anomalous results did not recur and the tests generally conformed with the expected null hypothesis distribution. It thus appeared that this process had produced a more homogeneous dataset which was used in the subsequent analyses.

In order to construct an exome-wide risk score the aim was to follow the approach implemented in SCOREASSOC and provide a weight for each variant based on its frequency and predicted function (Curtis 2012). Thus, subjects with more rare, functional variants would receive higher scores. A gene-wise risk score is derived as the sum of the variant-wise weights, each multiplied by the number of alleles of the variant which a given subject possesses. An exome-wide risk score can be derived as the weighted sum of gene-wise scores, with some genes being weighted more highly than others. Potentially such a model has a large number of parameters because a different weight can be assigned to each variant and to each gene.

Each variant was annotated using VEP, PolyPhen and SIFT (McLaren et al. 2016)(Adzhubei et al. 2013; Kumar et al. 2009). VEP produces annotations for 36 different possible types of variant. In order to reduce the parameter space, each variant type was characterised according to whether each of seven attributes was applicable to it, these attributes being: possibly having a non-coding effect through being in a regulatory region or intronic; in UTR; in coding region; nonsynonymous; loss of function; possibly or probably damaging according to PolyPhen; deleterious according to SIFT. Table 1 shows the list of VEP annotation types along with which attributes would be applicable to each. Each attribute would be assigned a weight and then the weight for a particular variant would consist of the sum of weights of its applicable attributes. A background weight for the attribute “any variant” would also be assigned, meaning that in all a total of eight attribute weights could be used to generate functional weights for all variants. So, for example, the weight for a variant annotated as 5’ UTR would be the sum of three weights for *any variant, possible non-coding effect* and UTR.

**Table 1.** Attributes ascribed to VEP annotation types. Each 1 indicates that the relevant attribute was assigned to the annotation. Three additional attributes were used. All variants were considered to have the “Any variant” attribute. Independent of their VEP annotation, variants could be classified as “Deleterious” by SIFT and variants could be classified as “Possibly or probably damaging” by PolyPhen. Thus each variant could be assigned up to eight annotations.

| VEP | Possible non-coding effect | UT R | Codin g | Non-synonymou s | LOF |
| --- | --- | --- | --- | --- | --- |
| NULL_CONSEQUENCE |  |  |  |  |  |
| intergenic_variant |  |  |  |  |  |
| feature_truncation | 1 |  |  |  |  |
| regulatory_region_variant | 1 |  |  |  |  |
| feature_elongation | 1 |  |  |  |  |
| regulatory_region_amplification | 1 |  |  |  |  |
| regulatory_region_ablation | 1 |  |  |  |  |
| TF_binding_site_variant | 1 |  |  |  |  |
| TFBS_amplification | 1 |  |  |  |  |
| TFBS_ablation | 1 |  |  |  |  |
| downstream_gene_variant | 1 |  |  |  |  |
| upstream_gene_variant | 1 |  |  |  |  |
| non_coding_transcript_variant | 1 |  |  |  |  |
| NMD_transcript_variant | 1 |  |  |  |  |
| intron_variant | 1 |  |  |  |  |
| non_coding_transcript_exon_variant | 1 |  |  |  |  |
| 3_prime_UTR_variant | 1 | 1 |  |  |  |
| 5_prime_UTR_variant | 1 | 1 |  |  |  |
| mature_miRNA_variant | 1 |  |  |  |  |
| coding_sequence_variant |  |  | 1 |  |  |
| synonymous_variant |  |  | 1 |  |  |
| stop_retained_variant |  |  | 1 |  |  |
| incomplete_terminal_codon_variant |  |  | 1 |  |  |
| splice_region_variant |  |  | 1 |  |  |
| protein_altering_variant |  |  | 1 | 1 |  |
| missense_variant |  |  | 1 | 1 |  |
| inframe_deletion |  |  | 1 | 1 |  |
| inframe_insertion |  |  | 1 | 1 |  |
| transcript_amplification |  |  | 1 | 1 |  |
| start_lost |  |  | 1 | 1 |  |
| stop_lost |  |  | 1 | 1 |  |
| frameshift_variant |  |  | 1 | 1 | 1 |
| stop_gained |  |  | 1 | 1 | 1 |
| splice_donor_variant |  |  | 1 | 1 | 1 |
| splice_acceptor_variant |  |  | 1 | 1 | 1 |
| transcript_ablation |  |  | 1 | 1 | 1 |

For a general application, the weight for each variant would also be multiplied by a factor based on its frequency, with rarer variants being given higher weight. However previous research has made it clear that there are no common variants with a substantial effect on risk of schizophrenia and hence it was decided to restrict attention to variants with MAF of 0.01 or less in either cases or controls. In these circumstances the weighting scheme based on frequency as implemented in SCOREASSOC would have had a negligible effect in terms of distinguishing between rare and extremely rare variants and so no frequency-based weighting was applied. Applying the above QC processes and allele frequency restriction yielded genotypes for 1,177,741 variants in 19,627 genes.

In principle, with improved knowledge about the genetic contribution to schizophrenia risk it would be possible to assign weights to individual genes. At present it is not clear which individual genes are involved or the magnitude of their associated risks. However previous work has proposed sets of genes which may be enriched for rare, functional variants in schizophrenia cases (Purcell et al. 2014; Curtis 2016; Genovese et al. 2016). The lists of genes for the gene sets tested for enrichment in the original analysis of this dataset are shown in Table 2. In addition to these, a set was created of which all genes were a member. Rather than assign a weight to each gene separately, a weight could be assigned to each gene set and then the weight for a gene could be defined as the sum of the weights of all the gene sets of which it was a member.

**Table 2.** Gene sets used in in the original analysis of this dataset which provides a full description of their derivation is in the online methods section (Genovese et al. 2016). The lists were obtained directly from the first author. The symbol used is the same as that used for the name of the file containing the list.

| Gene set | Symbol |
| --- | --- |
| OMIM intellectual disability (Hamosh et al. 2005) | *alid* |
| Expression specific to brain (Fagerberg et al. 2014) | *brain* |
| Bound by CELF4 (Wagnon et al. 2012) | *celf4* |
| Missense-constrained (Samocha et al. 2014) | *constrained* |
| Involved in developmental disorder (McRae et al. 2017) | *dd* |
| De novo variants in autism (Fromer et al. 2014) | *denovo.aut* |
| De novo variants in coronary heart disease (Fromer et al. 2014) | *denovo.chd* |
| De novo variants in epilepsy (Fromer et al. 2014) | *denovo.epi* |
| De novo duplications in ASD (Kirov et al. 2012) | *denovo.gain.asd* |
| De novo duplications in bipolar disorder (Kirov et al. 2012) | *denovo.gain.bd* |
| De novo duplications in schizophrenia (Kirov et al. 2012) | *denovo.gain.scz* |
| De novo variants in intellectual disability (Fromer et al. 2014) | *denovo.id* |
| De novo deletions in ASD (Kirov et al. 2012) | *denovo.loss.asd* |
| De novo deletions in bipolar disorder (Kirov et al. 2012) | *denovo.loss.bd* |
| De novo deletions in schizophrenia (Kirov et al. 2012) | *denovo.loss.scz* |
| De novo variants in schizophrenia (Fromer et al. 2014) | *denovo.scz* |
| Bound by FMRP (Darnell et al. 2011) | *fmrp* |
| Implicated by GWAS (Schizophrenia Working Group of the Psychiatric Genomics Consortium 2014) | *gwas* |
| Targets of microRNA-137 (Robinson et al. 2015) | *mir137* |
| Expression specific to neurons (Cahoy et al. 2008) | *neurons* |
| NMDAR and ARC complexes (Kirov et al. 2012) | *nmdarc* |
| Loss-of-function intolerant (Lek et al. 2015) | *pLI09* |
| PSD-95 (Bayés et al. 2011) | *psd95* |
| Bound by RBFOX 1 or 3 (Weyn-Vanhentenryck et al. 2014) | *rbfox13* |
| Bound by RBFOX 2 (Weyn-Vanhentenryck et al. 2014) | *rbfox2* |
| Synaptic (Pirooznia et al. 2012) | *synaptome* |
| Escape X-inactivation (Cotton et al. 2013) | *x.escape* |
| X-linked intellectual disability, Genetic Services Laboratories of the University of Chicago (Moeschler 2008; Géczy et al. 2009; Moeschler et al. 2006; Rauch et al. 2006) | *xlid.chicago* |
| X-linked intellectual disability, Greenwood Genetic Centre (Moeschler et al. 2006) | *xlid.gcc* |
| X-linked intellectual disability, OMIM (Hamosh et al. 2005) | *xlid.omim* |
| X-linked intellectual disability (combined) | *xlid* |

With this approach in mind, an overall risk score can be calculated as the product of a number of matrices, as follows:

A is a matrix which defines which attributes are possessed by each variant. It has columns equal to N_Var_, the number of variants, and rows equal to N_Attrib_, the number of attributes, here 8. A_ij_ is 1 or 0, depending on whether the *j*th variant has the *i*th attribute.

F is diagonal matrix with N_Var_ rows and columns. The diagonal elements consist of weights derived from the allele frequency so that variants with high MAF have a weight close to 1 and rare variants have a weight close to an arbitrarily chosen weighting factor, as implemented in SCOREASSOC and as described previously (Curtis 2012). As stated above, this weighting was not applied for the current analyses, equivalent to setting all diagonal elements to 1.

I is the indicator matrix which codes the subject genotype at each variant. It is a diagonal matrix with N_Var_ rows and columns and the diagonal elements consist of 0, 1 or 2 depending on how many copies of the minor allele of the variant the subject possesses. If a subject had an unknown genotype they would be assigned a value of 2xMAF.

G is a matrix with N_Var_ rows and number of columns equal to N_Gene_, the number of genes tested. G_ij_ is 1 if the *i*th variant is in the *j*th gene, with the other elements of the row being 0. Variants were extracted and dealt with one gene at a time and each variant was assigned to the gene for which it was extracted. Since for each gene all variants were extracted between the transcription start and end sites, a small number of variants in overlapping genes would have been extracted twice and would be dealt with as two different variants, each assigned to a different gene.

S is a matrix with N_Gene_ rows and number of columns equal to N_Set_, the number of gene sets used. S_ij_ is 1 if the *i*th gene is a member of the *j*th gene set and 0 otherwise. Since a gene can be a member of more than 1 set, there could be several 1 values in each row.

W_Att_ is a row vector with N_Att_ elements providing the weights for each attribute.

W_Set_ is column vector with N_Set_ elements providing the weights for each gene set.

Using this notation, the overall risk score R for a subject is given by:

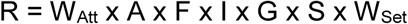

Only the values for elements of I differ between subjects.

In order to allow rapid recalculation of the risk score for different values of the weights for the gene sets and variant attributes, it is helpful to calculate for each subject an intermediate matrix D with N_Set_ columns and N_Att_ rows which contains a summary of aggregate scores by gene set and attribute so that we have:

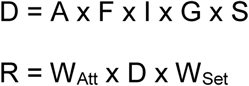

In order to implement this system in practice the following procedure was applied. VEP, SIFT and PolyPhen annotations were obtained for all the variants in the case-control VCF file. GENEVARASSOC was used to extract the genotypes for variants one gene at a time and used the annotations to provide a weight for each variant consisting of a binary number denoting the attributes which were applicable to that variant. (So that a variant with the second and third attributes would have a weight of 110 = 6.) Next, a custom-written program was used to produce aggregate attribute scores from the variant scores by decoding the weight to determine which attributes were applicable to each variant. At this stage, weighting for frequency could also have been applied. Using the above notation, this was equivalent to obtaining A x F x I for each subject and each gene. Finally, these attribute scores for each gene were combined into attribute scores for each gene set based on which genes were members of each set This resulted in a condensed dataset consisting of, for each subject, the aggregate scores for each attribute and gene set, denoted D above.

As described previously, a weighted burden test can be carried out by performing a two sample t test to compare the risk score, R, between cases and controls (Curtis 2012). In order to find a set of weights which best distinguishes cases from controls we can simply seek to maximise this t statistic. A program was written which would:

1. Read in the subject-wise scores aggregated by attribute and gene set along with a set of weights for attributes and gene sets;
2. calculate the t statistic;
3. maximise the t statistic over different values for the weights using Powell’s conjugate direction method, which does not require that a function be differentiable (Powell 1964).

Initially, maximisation of the t statistic was carried out for the whole dataset for all 8 attribute weights and 36 gene set weights in order to find the best-fitting values. For each weight a "1 t confidence interval” was then defined as the range of values which could be assigned to that weight, keeping all other weights fixed, which would yield a t statistic no less than the maximum t statistic minus 1.

To find a good-fitting minimal set of weights a step-wise procedure was followed. The weight for each attribute or gene set in turn was set to 0 and the t statistic was recalculated. If any produced a reduction in the t statistic to less than 1 below the original maximum the weight producing the smallest reduction was fixed at 0 and then the maximisation was repeated again over all the surviving weights.

In order to assess the statistical significance of the fitted risk scores, a five-way leave-one-out cross-validation procedure was used. Maximisation to find the best-fitting weights was carried out in a training four-fifths of the dataset and then risk scores were calculated using these weights in the remaining test fifth. In addition, the t statistic which would have been obtained in the entire sample using these weights was calculated. This was repeated five times. The risk scores from each test fifth were then standardised by subtracting the mean and dividing by the standard deviation and then all five were combined and a t test was performed on the standardised risk scores for the whole sample.

In order to assess the statistical significance of fitted risk scores derived from a minimal set of weights the step-wise process described above was carried for each four-fifths and then the weight obtained were used to calculate risk scores in the remaining fifth. Again, the combined, standardised risk scores obtained from the test subjects were then compared using a t test.

The ability of the standardised risk scores in the test subjects to distinguish cases from controls was by calculating the receiver operating characteristic curve using the pROC package (Robin et al. 2011).

The results are affected by the relative rather than absolute values of the weights, so in order to aid comparison of the results in the tables all the fitted weights were scaled so that the average magnitude for gene set weights and for the attribute weights would be 10.

## Results

Fitting all gene set and attribute weights produced a maximised t statistic of 9.5 with the fitted values shown in Table 3. As can be seen, many weights had a wide confidence interval which included 0 and hence could be taken not to materially contribute to the fit. Applying the stepwise procedure to retain only important weights resulted in the minimal set also shown in Table 3. This includes weights for 8 out of the 32 gene sets and 3 of the attributes and with this reduced parameter set it was possible to produce a maximised t statistic of 8.6. The attributes with positive weights in this model were *any variant, PolyPhen damaging* and *SIFT deleterious*. The weights for both *nonsynonymous* and *LOF* could be set to 0, presumably because variants with this consequence could be adequately weighted using the SIFT and PolyPhen attributes.

**Table 3.** Fitted weights chosen to maximise the t statistic for the risk score to distinguish cases from controls using all weights and using a minimal set of weights. The range indicates the values that the weight can take without reducing the t statistic to less than the maximum achieved minus 1. The minimal set of weights consists of those which are needed to produce a t statistic not less than the maximum minus 1.

|  | Fitted weights<br>(range) | Minimal fitted<br>weights<br>(range) |
| --- | --- | --- |
| Gene sets |  |  |
| *alid* | 10.8<br>(-44.1,71.1) | 0 |
| *brain* | -1.7<br>(-6.6,3.8) | 0 |
| *celf4* | -4.4<br>(-13.3,5.7) | 0 |
| *constrained* | 2.4<br>(-12.1,21.5) | 0 |
| *dd* | -44.1<br>(-111.2,9.4) | -15.3<br>(-24.8,-7.6) |
| *denovo.aut* | -0.1<br>(-4.2,5.2) | 0 |
| *denovo.chd* | 6.2<br>(-10.3,28.9) | 2.7<br>(0.4,5.4) |
| *denovo.epi* | 1.2<br>(-14.1,19.3) | 0 |
| *denovo.gain.asd* | -0.0<br>(-10.7,12.3) | 0 |
| *denovo.gain.bd* | 7.3<br>(-26.8,48.0) | 0 |
| *denovo.gain.scz* | -5.0<br>(-34.9,24.1) | 0 |
| *denovo.id* | 1.1<br>(-6.2,9.4) | 0 |
| *denovo.loss.asd* | 2.3<br>(-6.4,13.8) | 0 |
| *denovo.loss.bd* | -7.5<br>(-55.8,52.0) | 0 |
| *denovo.loss.scz* | 11.5<br>(-16.2,47.6) | 5.2<br>(1.1,9.9) |
| *denovo.scz* | 0.1<br>(-6.0,6.8) | 0 |
| *fmrp* | 0.5<br>(-6.2,9.2) | 0 |
| *gwas* | 26.5<br>(-13.6,84.4) | 13.5<br>(7.2,21.4) |
| *mir137* | 1.5<br>(-5.0,11.2) | 0 |
| *neurons* | 2.8<br>(-4.3,13.1) | 0 |
| *nmdarc* | 9.8<br>(-40.7,74.1) | 0 |
| *pLI09* | 3.6<br>(-3.3,15.3) | 2.2<br>(1.1,3.7) |
| *psd95* | -3.6<br>(-55.7,61.5) | 0 |
| *rbfox13* | -1.6<br>(-8.7,7.3) | 0 |
| *rbfox2* | 0.6<br>(-6.7,10.1) | 0 |
| *synaptome* | 3.9<br>(-5.0,17.8) | 1.7<br>(0.2,3.4) |
| *x.escape* | 40.4<br>(5.5,111.5) | 18.5<br>(12.6,26.8) |
| *xlid.chicago* | -70.1<br>(-185.8,42.8) | -20.9<br>(-34.8,-1.4) |
| *xlid.gcc* | 1.4<br>(-65.7,74.9) | 0 |
| *xlid.omim* | -17.9<br>(-110.6,77.0) | 0 |
| *xlid* | 30.1<br>(-23.6,95.6) | 0 |
| *any gene* | 0.4<br>(-1.1,2.9) | 0 |
| Variant attributes |  |  |
| *any variant* | 1.3<br>(-1.8,9.8) | 2.2<br>(1.3,3.5) |
| *non-coding effect* | 4.5<br>(-1.1,16.2) | 0 |
| *UTR* | -7.3<br>(-33.4,20.8) | 0 |
| *coding* | 0.5<br>(-4.0,10.0) | 0 |
| *nonsynonymous* | -0.8<br>(-9.2,18.4) | 0 |
| *LOF* | 17.9<br>(-52.2,99.2) | 0 |
| *PolyPhen damaging* | 15.5<br>(-9.2,87.6) | 7.5<br>(1.0,16.6) |
| *SIFT deleterious* | 32.1<br>(2.2,193.2) | 20.3<br>(11.2,35.4) |
| t statistic achieved | 9.5 | 8.6 |

The weights fitted for each of the five training sets are shown in Table 4. Each set consists of a different four fifths of the dataset and hence they overlap with each other and the fitted weights they yield are similar though not identical. When the fitted weights were used to calculate a t statistic in the whole sample, the different training sets produced values ranging from 8.4 to 9.2, showing that each set of weights represented a solution reasonably close to the best attainable. The scores for the test samples in each fifth not used for training were standardised and combined and then a t test was performed comparing scores in cases and controls. This produced a t statistic of 3.3 with a p value of 0.001, demonstrating that the risk scores which are produced are indeed associated with risk of schizophrenia and are not simply an artefact of the fitting process.

**Table 4.** Fitted weights chosen to maximise the t statistic in five training sets, each consisting of four fifths of the total sample. Also shown is the t statistic which is produced in the whole sample using that set of weights.

|  | Fitted weights<br>1 (range) | Fitted weights<br>2 (range) | Fitted weights<br>3 (range) | Fitted weights<br>4 (range) | Fitted weights<br>5 (range) |
| --- | --- | --- | --- | --- | --- |
| Gene sets |  |  |  |  |  |
| *alid* | 15.9<br>(-24.2,61.8) | -2.3<br>(-114.2,113.8) | 2.5<br>(-48.2,53.8) | -4.8<br>(-75.2,64.3) | 23.5<br>(-27.0,82.8) |
| *brain* | -1.0<br>(-4.9,3.3) | -3.9<br>(-10.2,2.2) | -0.8<br>(-5.1,4.1) | -1.8<br>(-6.1,2.5) | -1.4<br>(-6.7,4.9) |
| *celf4* | -3.2<br>(-10.1,4.5) | 0.5<br>(-14.4,23.0) | -4.6<br>(-12.5,3.9) | -1.4<br>(-10.9,10.5) | -6.8<br>(-16.5,2.5) |
| *constrained* | -0.7<br>(-11.6,11.8) | 6.6<br>(-20.5,46.7) | 0.8<br>(-12.8,17.4) | 0.8<br>(-16.3,21.7) | 3.6<br>(-10.7,21.7) |
| *dd* | -28.8<br>(-76.5,10.3) | -42.8<br>(-154.7,63.7) | -36.7<br>(-97.8,11.8) | -48.3<br>(-128.4,15.6) | -48.1<br>(-116.4,7.6) |
| *denovo.aut* | -1.3<br>(-4.6,2.4) | 1.4<br>(-4.1,8.1) | -0.1<br>(-3.8,4.6) | 3.2<br>(-0.5,7.9) | -0.4<br>(-4.9,5.3) |
| *denovo.chd* | 4.6<br>(-8.1,20.7) | 9.2<br>(-13.3,39.7) | 5.7<br>(-9.2,26.2) | 7.5<br>(-9.0,30.2) | 1.2<br>(-16.5,22.1) |
| *denovo.epi* | -2.9<br>(-15.2,9.4) | 4.3<br>(-15.2,30.6) | 1.5<br>(-12.6,17.8) | 0.8<br>(-15.3,19.5) | 4.2<br>(-11.7,24.1) |
| *denovo.gain.asd* | 1.1<br>(-7.0,10.8) | -1.0<br>(-20.5,21.5) | 0.5<br>(-8.6,11.4) | -1.0<br>(-13.1,12.9) | -1.2<br>(-12.9,11.3) |
| *denovo.gain.bd* | 8.1<br>(-18.2,40.2) | 9.9<br>(-44.8,74.0) | 5.9<br>(-25.2,42.2) | 14.2<br>(-15.3,54.1) | -13.9<br>(-56.4,23.8) |
| *denovo.gain.scz* | -7.8<br>(-31.9,13.6) | -13.4<br>(-69.3,40.5) | -1.0<br>(-26.4,24.2) | 3.3<br>(-24.2,34.6) | -1.1<br>(-32.8,29.8) |
| *denovo.id* | 0.8<br>(-5.1,7.3) | -3.5<br>(-11.6,4.6) | 1.2<br>(-5.7,8.9) | 0.5<br>(-5.6,6.8) | 4.6<br>(-3.7,14.7) |
| *denovo.loss.asd* | 1.7<br>(-5.2,10.4) | 2.1<br>(-12.4,21.4) | 1.5<br>(-6.0,11.0) | 1.5<br>(-7.4,13.0) | 1.6<br>(-8.5,13.9) |
| *denovo.loss.bd* | -1.9<br>(-41.4,44.4) | 0.5<br>(-92.0,117.0) | -3.8<br>(-44.0,48.0) | 8.7<br>(-43.8,84.6) | -14.8<br>(-63.9,38.9) |
| *denovo.loss.scz* | 5.3<br>(-17.6,33.0) | 12.0<br>(-36.1,70.9) | 13.6<br>(-9.3,47.9) | 13.8<br>(-16.9,56.3) | 8.0<br>(-20.3,41.5) |
| *denovo.scz* | 0.5<br>(-4.2,5.4) | -0.5<br>(-7.8,6.6) | 0.2<br>(-5.5,6.3) | -2.8<br>(-8.1,2.1) | 0.2<br>(-6.7,8.1) |
| *fmrp* | -0.1<br>(-5.4,6.6) | 5.7<br>(-2.8,16.0) | -0.1<br>(-6.4,7.8) | 0.0<br>(-6.3,7.3) | -2.9<br>(-10.2,6.0) |
| *gwas* | 20.1<br>(-11.2,64.4) | 51.8<br>(-13.9,158.5) | 18.9<br>(-15.4,66.4) | 9.4<br>(-30.5,55.7) | 27.0<br>(-15.5,86.9) |
| *mir137* | 0.7<br>(-4.6,7.6) | 1.9<br>(-9.2,20.2) | 1.7<br>(-4.0,10.0) | 1.8<br>(-5.3,12.1) | 0.7<br>(-6.2,9.8) |
| *neurons* | 4.4<br>(-1.1,12.7) | -0.2<br>(-12.9,17.1) | 2.2<br>(-3.9,10.9) | 0.4<br>(-7.5,10.3) | 2.9<br>(-4.4,12.6) |
| *nmdarc* | 9.0<br>(-29.9,58.3) | 32.3<br>(-69.4,181.6) | 2.9<br>(-39.8,56.0) | 27.8<br>(-27.9,107.7) | 0.6<br>(-50.5,59.5) |
| *pLI09* | 2.5<br>(-2.6,10.2) | 8.6<br>(-3.9,36.5) | 3.5<br>(-2.6,14.0) | 5.1<br>(-2.8,18.6) | 2.8<br>(-4.0,12.8) |
| *psd95* | 6.4<br>(-35.9,61.5) | 11.9<br>(-92.2,155.1) | -3.0<br>(-45.9,51.3) | -2.3<br>(-63.4,75.0) | -10.8<br>(-64.3,46.3) |
| *rbfox13* | -0.8<br>(-6.3,5.7) | -3.2<br>(-15.3,15.3) | 1.2<br>(-5.1,9.1) | -1.9<br>(-9.8,8.0) | -2.4<br>(-9.9,5.9) |
| *rbfox2* | -1.0<br>(-6.7,5.7) | 6.6<br>(-5.7,27.5) | -2.4<br>(-8.9,5.5) | 2.3<br>(-5.6,13.0) | 1.0<br>(-6.6,9.9) |
| *synaptome* | 2.7<br>(-4.0,12.6) | 0.9<br>(-15.4,25.4) | 4.8<br>(-2.9,17.9) | 1.0<br>(-8.9,14.3) | 3.7<br>(-5.2,16.6) |
| *x.escape* | 30.6<br>(1.5,88.5) | 57.8<br>(-3.9,175.9) | 28.5<br>(-0.2,80.2) | 36.5<br>(2.4,106.4) | 37.2<br>(4.7,97.5) |
| *xlid.chicago* | -74.0<br>(-166.3,13.1) | -15.0<br>(-251.7,223.7) | -43.6<br>(-139.9,55.3) | -71.5<br>(-212.7,57.0) | -52.9<br>(-175.4,48.0) |
| *xlid.gcc* | -10.9<br>(-66.2,48.2) | 4.7<br>(-133.6,168.6) | -23.5<br>(-76.6,42.2) | 10.7<br>(-73.8,100.8) | -2.0<br>(-69.3,67.5) |
| *xlid.omim* | -23.5<br>(-100.8,58.0) | 4.4<br>(-139.1,168.7) | -44.4<br>(-120.7,38.3) | -20.8<br>(-129.2,86.0) | -11.9<br>(-109.2,85.8) |
| *xlid* | 47.4<br>(2.5,101.9) | -0.6<br>(-103.8,123.8) | 58.9<br>(15.2,116.8) | 14.2<br>(-53.5,88.1) | 25.1<br>(-29.0,89.6) |
| *any gene* | 0.4<br>(-0.7,2.1) | -0.5<br>(-2.8,2.8) | 0.2<br>(-1.1,2.3) | -0.2<br>(-1.7,1.9) | 1.3<br>(-0.4,4.2) |
| Variant attributes |  |  |  |  |  |
| *any variant* | 1.8<br>(-1.3,11.1) | 1.1<br>(-2.2,17.0) | 0.6<br>(-2.5,7.3) | -0.0<br>(-2.1,5.8) | 1.8<br>(-0.9,8.1) |
| *non-coding effect* | 4.8<br>(-0.9,16.7) | 1.3<br>(-4.4,11.6) | 4.7<br>(-1.0,15.4) | 2.0<br>(-1.9,8.5) | 5.1<br>(-0.4,17.8) |
| *UTR* | -13.3<br>(-38.2,11.2) | -5.4<br>(-36.7,29.7) | -4.9<br>(-32.6,26.2) | 1.5<br>(-15.4,22.4) | -5.3<br>(-29.4,21.6) |
| *coding* | 0.1<br>(-4.4,9.8) | 3.5<br>(-1.8,42.5) | 0.1<br>(-4.4,8.2) | 2.4<br>(-0.7,11.1) | -1.8<br>(-5.7,4.7) |
| *nonsynonymous* | 0.0<br>(-8.5,18.3) | 3.7<br>(-6.8,74.6) | -0.8<br>(-9.2,16.1) | -0.5<br>(-6.6,14.8) | -0.3<br>(-7.8,13.0) |
| *LOF* | 22.7<br>(-42.0,104.8) | 32.5<br>(-44.8,138.6) | 14.3<br>(-65.0,103.6) | 47.0<br>(-3.1,135.9) | -21.3<br>(-92.8,40.6) |
| *PolyPhen*<br>*damaging* | 0.9<br>(-20.8,43.4) | 10.8<br>(-15.9,74.9) | 16.2<br>(-9.9,113.1) | 11.3<br>(-5.6,55.4) | 26.4<br>(1.4,131.7) |
| *SIFT deleterious* | 36.4<br>(9.9,160.3) | 21.7<br>(-9.2,101.2) | 38.4<br>(5.3,576.3) | 15.1<br>(-4.2,69.0) | 18.1<br>(-7.8,101.8) |
| t statistic in<br>whole dataset | 8.8 | 8.4 | 9.2 | 8.8 | 8.7 |

The fitted weights produced by applying the step-wise procedure to each training set are shown in Table 5. It can be seen that there is considerable variation in the parameters retained. However the t statistics obtained for the whole sample using these weights varied between 7.4 and 8.3, showing that the different combinations of parameters selected were all able to produce risk scores which differed between cases and controls. When the standardised scores from the cases and controls not used for training were compared, the results were significant with a t statistic of 3.4 and a p value of 0.0006. Using either the full set of parameters or the minimal set, the ability of the risk score to distinguish cases from controls was extremely modest, with an area under the curve of only 0.52 in both situations.

**Table 5.** Minimal set of weights for each training set. The weights are chosen to produce a t statistic not less than the maximum produced in the training set minus 1. Also shown is the t statistic produced in the whole dataset using the minimal set of weights.

|  | Minimal weights 1<br>(range) | Minimal weights 2<br>(range) | Minimal weights 3<br>(range) | Minimal weights 4<br>(range) | Minimal weights 5<br>(range) |
| --- | --- | --- | --- | --- | --- |
| Gene sets |  |  |  |  |  |
| *alid* | 4.3<br>(1.4,7.4) | 0 | 0 | 0 | 10.8<br>(3.9,18.7) |
| *brain* | 0 | -1.6<br>(-2.7,-0.5) | 0 | 0 | 0 |
| *celf4* | -1.1<br>(-1.6,-0.6) | 0 | 0 | 0 | -3.2<br>(-4.5,-1.9) |
| *constrained* | 0 | 0 | 0 | 0 | 0 |
| *dd* | -8.2<br>(-11.3,-5.3) | 0 | -11.1<br>(-14.6,-7.8) | -19.0<br>(-29.9,-8.9) | -26.1<br>(-35.4,-18.8) |
| *denovo.aut* | 0 | 0 | 0 | 1.5<br>(1.0,2.0) | 0 |
| *denovo.chd* | 0 | 0 | 0 | 0 | 0 |
| *denovo.epi* | 0 | 3.7<br>(0.4,7.8) | 0 | 0 | 0 |
| *denovo.gain.asd* | 0 | 0 | 0 | 0 | 0 |
| *denovo.gain.bd* | 0 | 0 | 0 | 8.6<br>(3.9,13.9) | 0 |
| *denovo.gain.scz* | 0 | 0 | 0 | 0 | 0 |
| *denovo.id* | 0 | -1.6<br>(-3.1,-0.1) | 0 | 0 | 0 |
| *denovo.loss.asd* | 0 | 0 | 0 | 0 | 0 |
| *denovo.loss.bd* | 0 | 0 | 0 | 0 | 0 |
| *denovo.loss.scz* | 0 | 0 | 4.6<br>(2.9,6.3) | 10.1<br>(4.8,16.0) | 0 |
| *denovo.scz* | 0 | 0 | 0 | -2.0<br>(-2.7,-1.5) | 0 |
| *fmrp* | 0 | 3.7<br>(2.0,5.6) | 0 | 0 | 0 |
| *gwas* | 7.3<br>(4.6,10.4) | 25.8<br>(11.3,44.3) | 7.1<br>(4.6,9.8) | 0 | 14.5<br>(8.0,21.6) |
| *mir137* | 0 | 0 | 0 | 0 | 0 |
| *neurons* | 1.2<br>(0.7,1.7) | 0 | 0 | 0 | 0 |
| *nmdarc* | 0 | 0 | 0 | 18.6<br>(9.1,29.5) | 0 |
| *pLI09* | 0.9<br>(0.4,1.4) | 4.0<br>(1.5,7.7) | 1.1<br>(0.6,1.6) | 3.2<br>(1.7,4.9) | 1.6<br>(0.6,2.8) |
| *psd95* | 0 | 0 | 0 | 0 | 0 |
| *rbfox13* | 0 | 0 | 0 | 0 | 0 |
| *rbfox2* | 0 | 0 | 0 | 0 | 0 |
| *synaptome* | 0 | 0 | 1.6<br>(1.1,2.3) | 0 | 2.0<br>(0.7,3.5) |
| *x.escape* | 10.2<br>(7.5,13.5) | 29.5<br>(15.6,49.6) | 8.0<br>(5.8,10.4) | 17.0<br>(11.1,24.5) | 21.0<br>(14.7,27.7) |
| *xlid.chicago* | -17.4<br>(-23.3,-11.7) | 0 | 0 | 0 | 0 |
| *xlid.gcc* | -14.2<br>(-18.2,-10.1) | 0 | -14.6<br>(-18.1,-10.9) | 0 | 0 |
| *xlid.omim* | -18.2<br>(-23.7,-12.1) | 0 | -19.9<br>(-24.6,-15.2) | 0 | 0 |
| *xlid* | 27.0<br>(23.5,30.7) | 0 | 22.0<br>(19.1,25.1) | 0 | 0 |
| *any gene* | 0 | 0 | 0 | 0 | 0.7<br>(0.4,1.0) |
| Variant attributes |  |  |  |  |  |
| *any variant* | 1.7<br>(0.8,3.0) | 3.2<br>(1.3,7.7) | 0 | 0 | 0 |
| *non-coding effect* | 3.6<br>(2.1,5.5) | 0 | 2.6<br>(1.9,3.5) | 0 | 4.0<br>(3.0,5.5) |
| *UTR* | 0 | 0 | 0 | 0 | 0 |
| *coding* | 0 | 0 | 0 | 1.1<br>(0.6,1.8) | 0 |
| *nonsynonymous* | 0 | 6.7<br>(0.8,21.0) | 0 | 0 | 0 |
| *LOF* | 0 | 0 | 0 | 21.9<br>(14.8,31.0) | 0 |
| *PolyPhen damaging* | 0 | 0 | 9.8<br>(5.8,15.2) | 0 | 12.8<br>(7.9,20.1) |
| *SIFT deleterious* | 24.7<br>(18.0,33.8) | 20.2<br>(5.5,41.7) | 17.6<br>(12.3,25.5) | 7.0<br>(4.5,10.5) | 13.2<br>(7.3,20.7) |
| t statistic in whole dataset | 8.2 | 7.4 | 8.2 | 8 | 8.3 |

Thus, the minimal parameter sets found by the stepwise procedure result in risk scores which differ between cases from controls to a similar extent to the full set. The *SIFT deleterious* attribute is given a strong positive weight by all five training sets while *PolyPhen damaging* is used in two and *LOF* in one. The attribute for *non-coding effect* has a small positive weight in three of the training sets. With respect to gene sets, in all training sets *x.excape* is given a large positive weight and *pLI09* a small positive weight. These are the sets of genes which escape X inactivation and genes which are LOF intolerant. In four out of five training sets genes which are close to GWAS hits are given a strongly positive weight. Some gene sets are given negative weights. In four of the training sets *dd*, genes associated with developmental disability, is given a strongly negative weight and in two training sets the combined X-linked disability is given a positive weight but subsets of X-linked disability genes are weighted negatively. One way to interpret such findings is to view negative weights as encoding a “but not if “ relationship. For example, “escapes X inactivation but not if associated with developmental disability”. Of note is that there was no consistent retention of any of the gene sets which might be viewed as more specifically implicated in schizophrenia, such a de novo variants, or related to biology, such as neuronal, post-synaptic density or NMDA receptor genes.

## Discussion

The method presented here represents a first attempt to combine information from exome-wide variants into a single risk score. The association of the score with the trait in test subjects is statistically significant although with minimal effect size. One possible explanation for this is that the gene sets used provide a poor categorisation of which genes do and do not influence risk of schizophrenia. If this is the case then one would hope that performance of the method would improve as more knowledge is accumulated to lead to better definition of risk genes. However an alternative explanation would be that some of the gene sets do indeed consist mostly of genes influencing risk but that the variation within these genes is so widespread that only a small minority of variants have an effect and that the scheme used here to categorise variant effects is unable to distinguish them. Again, as additional knowledge emerges it might be possible to devise improved classification schemes which would feed into an improved weighting system.

The approach presented does provide a framework to systematically explore different kinds of contribution to risk. Given the complexity of the genetic architecture of schizophrenia, the sample sizes used here are too small for definite conclusions to be drawn but the results do illustrate the kind of inferences that could be made. For example, the results suggest that the SIFT prediction makes an important contribution to risk score but that other classifications do not provide much additional information. Likewise, the results suggest that genes which are loss of function intolerant, escape X inactivation or are implicated by a GWAS may be relevant to risk. However once these factors are taken into account the classifications which were chosen to reflect biological function do not appear to improve performance. The fact that some intellectual disability gene sets were given positive weights while others, including the genes for developmental disorder, were given negative weights hints at the notion that a subset of these genes influence schizophrenia risk and it is possible that one could use the risk score to explore this further. In general, the weighting of gene sets and variant attributes allows for a formal method to produce a summary risk score from all exome variants and to systematically explore the performance of different weighting schemes.

A risk score from exome variants could be combined with a polygenic risk score from common SNPs. It could also be combined with risk scores derived from identified rare variants which have been shown to have a major effect on risk, such as specific copy number variants and gene mutations (Raychaudhuri et al. 2010; Singh et al. 2016). A recent study of autism has demonstrated that in subjects who possess rare variants having major effects on risk for autism, common variants can increase this risk further (Weiner et al. 2017). Likewise, environmental factors could be incorporated to provide an overall assessment of disease risk.

The method as presented assumes that the cases and controls are drawn from the same population and does not include population principal components as covariates. Because it does not study individual variants but types of variant it may be relatively robust to differences in allele frequencies between sub-populations. On the other hand, if there were among cases an over-representation of a sub-population in which there was a higher frequency in general of rare variants then this would produce false positive results. Hence it seemed important to exclude all subjects which appeared to have a substantial contribution of Finnish ancestry since otherwise the excess of Finnish alleles among cases would have been problematic. Given that the contribution to risk score seemed to be confined to particular gene sets and variant types, it seems that this procedure did result in an acceptably homogeneous dataset.

The measure chosen to distinguish cases from controls was a weighted sum which could be used to obtain a t statistic. Other measures might be used, for example a log odds ratio which would fit into a logistic regression framework. The t statistic is very quick to calculate, which makes it attractive to use in the context of a maximisation process, and the risk score is simply a measure which increases with higher risk but which is not intended to provide a direct estimate of the actual quantitative risk of developing disease. The parameterisation of the model assumes that each type of variant affects each type of gene set equally. However more complex models could be developed, for example that loss of function variants in one set of genes increased risk but that for a different set of genes regulatory variants tended to be more important. Such models could be explored through machine learning techniques. Once a model had been developed using the general gene sets and variant types described above, it would be possible to try adding in additional genes or more specific gene sets in a systematic way in an effort to discover if any produced a significant improvement in the ability of the score to distinguish cases from controls. Such investigations will be the subject of subsequent work.

The scheme proposed here represents a starting point for a method to summarise the genetic risk contribution of variation at the level of the whole exome. As it stands, it is able to produce risk scores which are significantly different between schizophrenia cases and controls and hopefully its performance could be improved with information from additional datasets, refinement of gene sets and with further modifications to the procedure.

## Software availability

Source code of programs to calculate and optimise risk scores will be available from https://github.com/davenomiddlenamecurtis.

## Acknowledgements

The datasets used for the analysis described in this manuscript were obtained from dbGaP at http://www.ncbi.nlm.nih.gov/gap through dbGaP accession number phs000473.v2.p2. Samples used for data analysis were provided by the Swedish Cohort Collection supported by the NIMH grant R01MH077139, the Sylvan C. Herman Foundation, the Stanley Medical Research Institute and The Swedish Research Council (grants 2009–4959 and 2011–4659). Support for the exome sequencing was provided by the NIMH Grand Opportunity grant RCMH089905, the Sylvan C. Herman Foundation, a grant from the Stanley Medical Research Institute and multiple gifts to the Stanley Center for Psychiatric Research at the Broad Institute of MIT and Harvard.

